# Examining the interaction between TGFBR3 and ESRRA in ovarian cancer prognosis

**DOI:** 10.1101/080101

**Authors:** Alun Passey, Robert Brown, Charlotte S. Wilhelm-Benartzi

**Author notes:** To whom correspondence should be addressed: Dr Charlotte Wilhelm-Benartzi 3rd Floor Radiotherapy Building, Hammersmith Hospital Campus, Du Cane Road, London, W12 0NN.

## Abstract

The authors found that ESRRA when co-expressed at high levels with TGFβR3 may be prognostic for serous ovarian cancer overall survival in data from the Cancer Genome Atlas; however, this result was not validated in further datasets. A cell line was also identified in this study for future functional investigation of interactions between ESRRA and TGFβR3 in the context of oestrogen signaling in order to further elucidate their potential roles as prognostic biomarkers for serous ovarian cancer.

**FUNDING SUPPORT:** This work was supported by Cancer Research UK program A6689 (RB and CSWB) and the Medical Research Council (AP)

**CONFLICT OF INTEREST DISCLOSURES:** The authors have declared no conflicts of interest.

**AUTHOR CONTRIBUTIONS:** CSWB designed the study. AP, CSWB, and RB developed the methodology. AP and CSWB collected the data. AP, CSWB, and RB wrote the manuscript. CSWB is responsible for the overall content.

## INTRODUCTION

Ovarian cancer is the fourth most common cause of cancer-related death in women. Despite molecular and genetic advances, overall survival rates have not changed significantly in the past 30 years. New therapeutic targets and treatment approaches are needed to increase the 5 year survival rate from 40%.

Epidemiological studies have revealed that ovulation increases the risk of ovarian cancer, as exemplified by the increased risk seen with null parity and the decreased risk with oral contraceptive use (1); however, it is unknown what the impact of ovulation itself is on the prognosis of ovarian cancer patients. Ovulation is regulated by certain hormones, including Estrogen and Progesterone, and hormones such as these ovulation-related hormones have been shown to modulate the immune response in both humans and in animal models (2-3). It is further likely that the immune response itself plays an important role in cancer prognosis with immunotherapies currently being developed for many cancers (4).

The authors hypothesised that hormones have the ability to modulate the immune response by inducing changes in gene expression, potentially impacting patient prognosis and aimed to interrogate that hypothesis in this work. A further aim was to identify cell lines appropriate for future functional studies aiming to investigate the possible interaction between ovulation and immune related genes found above, oestrogen signalling and prognosis in serous ovarian cancer.

## METHODS

### Patient samples

The discovery Affymetrix HGU133A expression microarray and Illumina HumanMethylation27 Beadchip datasets for ovarian serous tumours were obtained from the publicly available TCGA data portal (5). The analysis was limited to tumours with methylation, expression and survival data, in total 308 tumours were included in the study. The first expression and methylation validation data was derived from the International Cancer Genome Consortium as described in (6). Briefly, 75 primary serous ovarian cancer tumour samples were included having RNA Seq expression data, Illumina Infinium HumanMethylation450 Beadchip data and with survival data. Another validation set for the expression results, included 256 serous ovarian cancer tumours having survival and Affymetrix HGU133A expression microarray data, as described in (7). The final methylation validation set was the Hammersmith Illumina HumanMethylation27 Beadchip dataset and included 71 serous ovarian cancer tumours with survival data, as described in (8).

### Cell culture

PEA1, PEA2, PEO1, PEO4, PEO14, PEO23, HEYA8, IGROV1, SKOV3, A2780 and CP70 cells were cultured in RPMI-1640/10% foetal bovine serum/5%L-glutamine and incubated at 37°C in 10% CO2, with the exception of OSE-C2 which were incubated at 33°C in 10% CO2.

### Bisulfite Modification and PCR

DNA extraction from cultured cells was performed using QIAamp® DNA Mini kit (Qiagen) according to manufacturer’s instructions. DNA from tumour and ascites pairs was obtained lyophilised from Prof. David ICGC. The DNA was quantified and purity determined (Nanodrop 1000, Thermo Scientific). 500ng of DNA was bisulfite converted using the EZ-DNA MethylationTM Kit D5001 (Zymo Research Corporation) according to manufacturer’s instructions. PCR was conducted using FastStart Taq DNA Polymerase kit (Roche) according to manufacturer’s instructions with forward and reverse primers at 0.8µM. All genes were amplified with the following protocol: initial denaturation 6 minutes at 95°C, denaturation 30 seconds at 95°C, annealing 30 seconds, extension 30 seconds at 72°C and final extension 5 minutes at 72°C. Denaturation, annealing and extension were repeated for 45 cycles. Annealing temperatures were 58°C for all genes with the exception of ESRRA which was 50°C. Products were run on a 2% agarose gel to ensure amplification prior to pyrosequencing.

### Pyrosequencing

Pyrosequencing was conducted using the Pyromark MD platform according to manufacturer’s instructions. 5µl of product was used per product plate well for all genes. Sequencing primers were used at 0.039µM. Quantification of CpG methylation was performed using Pyro Q-CpG software, data not meeting quality control requirements was excluded from further analysis. Pyrosequencing primers (including sequencing primers) were designed using Pyromark Assay Design 2.0 software with UCSC website sequences.

### Real-time PCR

Cell lysis and RNA extractions were performed using RNeasy (Qiagen) kit according to manufacturer’s instructions. The RNA was quantified and purity determined (Nanodrop 1000, Thermo Scientific). 2µg of RNA was reverse transcribed using RevertAid First Strand cDNA Synthesis Kit (Thermo Scientific) according to manufacturer’s instructions. Primers were designed using Ensembl gene sequences and the Primer3 online resource. Forward and reverse primers were designed to anneal in different exons. RT-PCR was conducted using iQTM SYBR® Green Supermix (BIO-RAD) according to the manufacturer’s instructions using 0.39µl of cDNA per well and primers at 0.4µM. The platform used was the C1000TM Thermal Cycler: CFX96TM Real-Time System (BIO-RAD). Expression was measured as an average corrected CT relative to that of GAPDH. Three technical replicates were taken for all measurements.

### Microarray processing

The expression microarray data have been pre-processed and normalized across the samples in the TCGA, ICGC and Tothill datasets as per (5-7). The methylation data have been summarized as β value (M/[M+U]), where M is the signal at the target CpG site from the methylation bead and U is the signals from the control bead and have been processed as per (5-6, 8-9) The poor quality probes have been excluded by TCGA (http://tcga-data.nci.nih.gov/docs/TCGA_Data_Primer.pdf and 5).

### Statistical analysis

Statistical analysis was performed using R (version 3.0.1) following the flowchart shown in Figure 1. The target genes were each split at the medians of each individual dataset for both the methylation and expression data and combined to create the four groups of interest. The association with overall survival (OS) was measured for expression and methylation separately using a multivariable Cox proportional hazard model adjusted for age at initial diagnosis, FIGO stage and batch (except for Tothill which did not have batch information); this association was not confounded by grade or surgical debulking across all datasets as checked in sensitivity analyses (data not shown). Descriptive Kaplan Meier curves are also shown and include the logrank p-value in their title to compare the survival distributions of the four combinations. Multiple comparisons were adjusted for using the false discovery rate (10).

**Figure 1.**
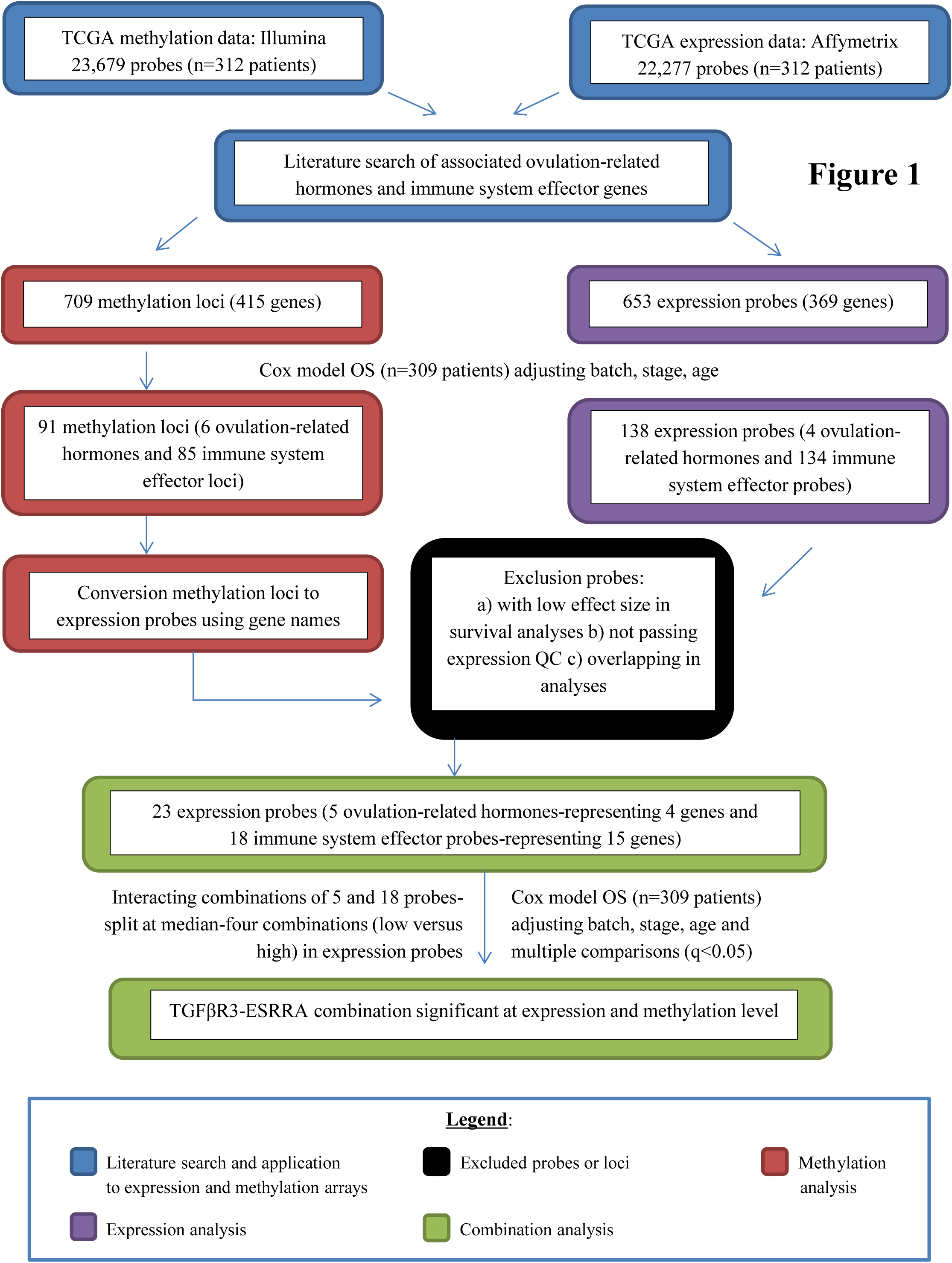
Flowchart of TCGA analysis.

## RESULTS

Biostatistical interrogation of The Cancer Genome Atlas (TCGA) clinical dataset revealed that high levels of expression (above the median) of one or both ESRRA and TGFβR3 are associated with poor OS in serous ovarian cancer patients. Correspondingly, low methylation levels of promoter-associated CpG sites of these gene pairs were associated with poor patient outcome; in the expected direction given that the methylation loci of interest were in the gene promoter region. Specifically, ovarian cancer patients with high levels of one or both receptors had an increased risk of death of at least two fold as compared to those patients with low levels of one or both receptors, after adjustment for potential confounders, multiple comparisons, and after running sensitivity analyses on grade and residual disease as shown in Table 1 and Figure 2a) and 2b). The interaction p-value was significant for expression of ESRRA and TGFβR3 in the TCGA dataset (Table 1).

**Figure 2.**
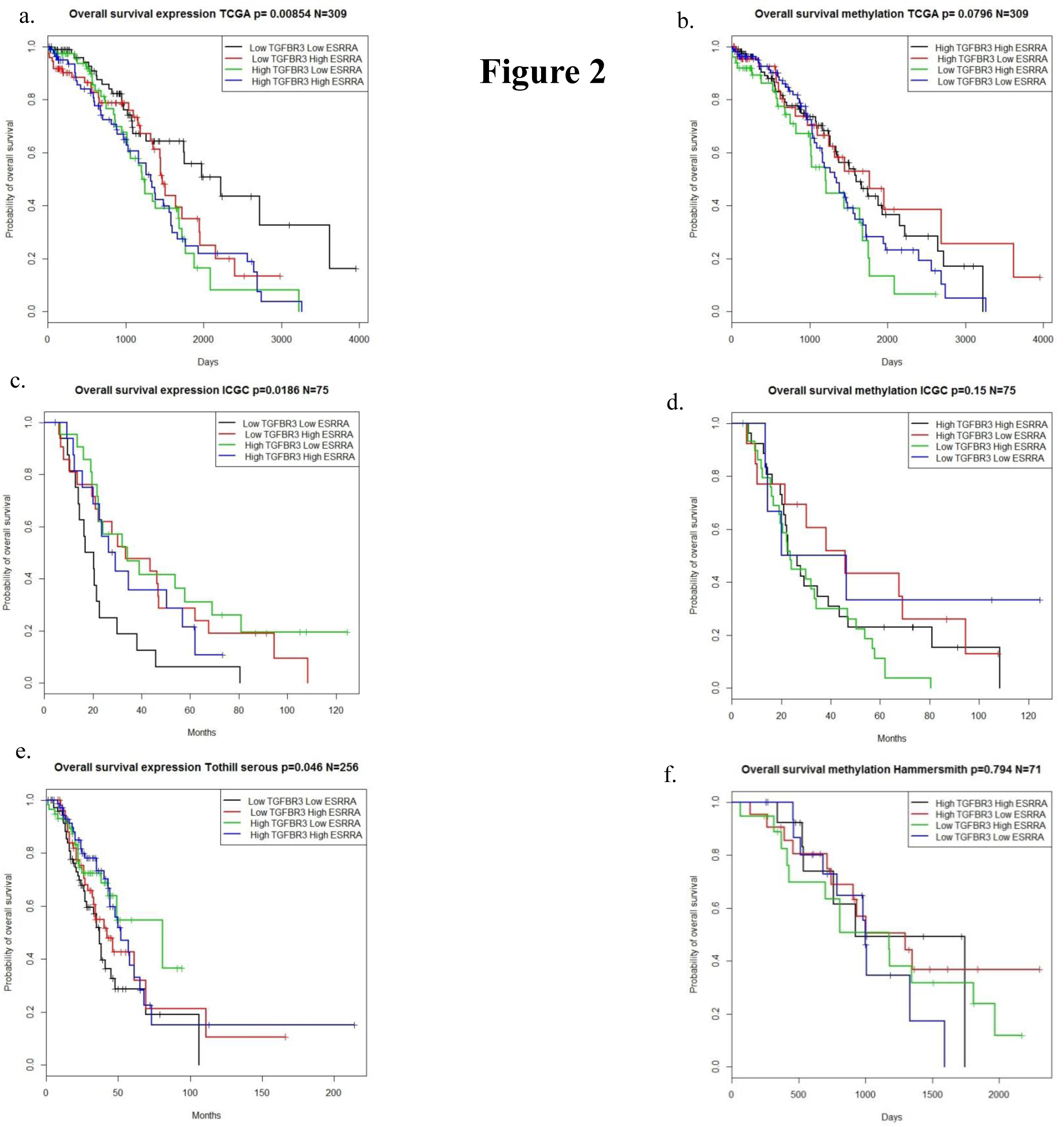
a. Kaplan Meier curve overall survival expression TCGA b. Kaplan Meier curve overall survival methylation TCGA c. Kaplan Meier curve overall survival expression ICGC d. Kaplan Meier curve overall survival methylation ICGC e. Kaplan Meier curve overall survival expression Tothill serous f. Kaplan Meier curve overall survival methylation Hammersmith.

**Table 1.**
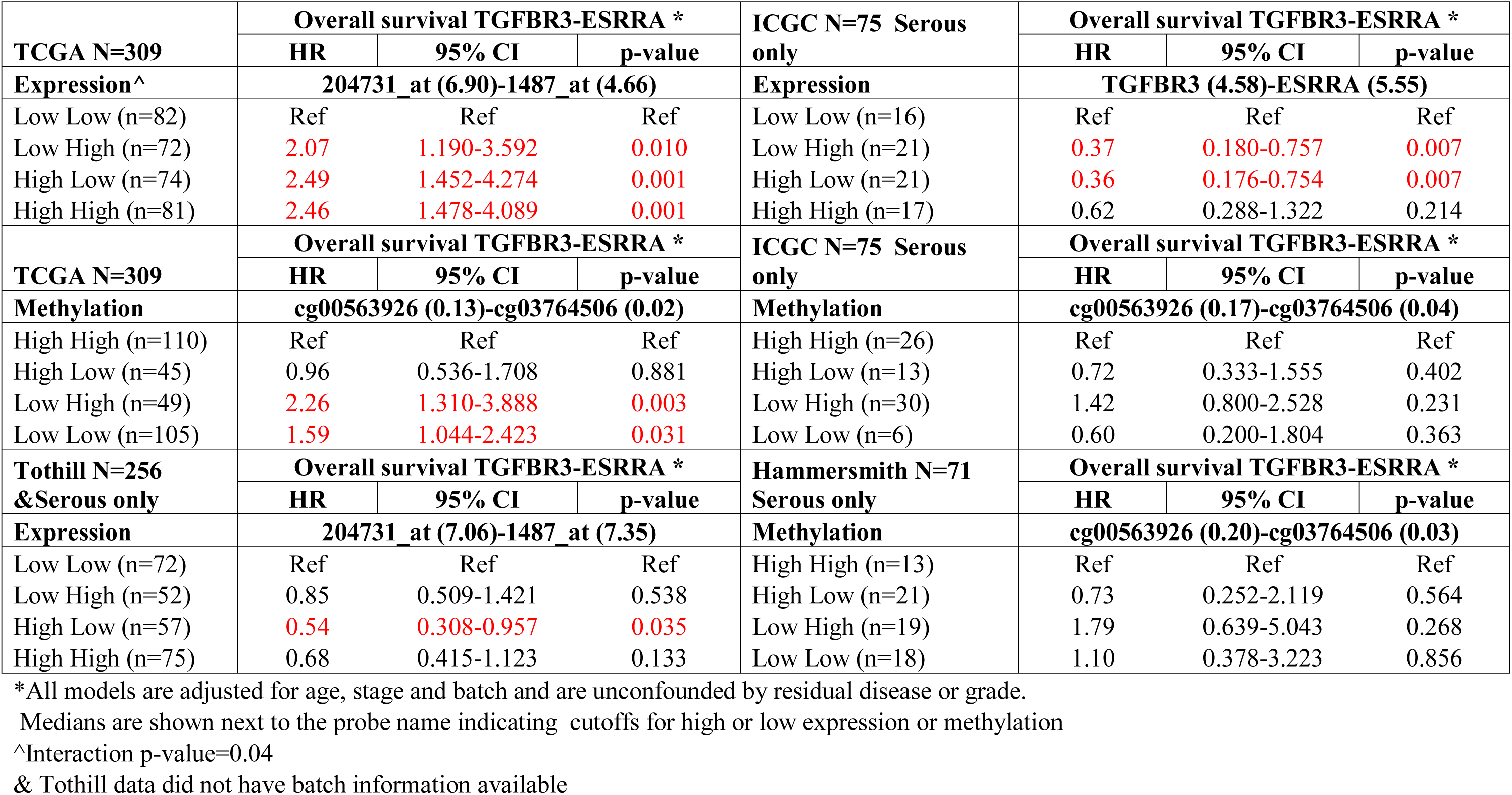
Statistical validation of TCGA expression and methylation results in ICGC, Tothill, and Hammersmith data.

Validation of these data was attempted in three datasets namely in the ICGC expression and methylation data (Table 1 and Figure 2c and 2d), in the Tothill expression data (Table 1 and Figure 2e) and in the Hammersmith methylation data (Table 1 and Figure 2f). Statistical analysis followed the same procedure as TCGA (Figure 1) with the probes or genes of interest extracted from the expression and methylation data and the four groups being created using the medians of each individual dataset. Potential confounders were adjusted for as well as sensitivity analyses run as per the TCGA analysis (Figure 1). Kaplan Meier curves were also constructed in Figure 2 for each dataset. Results were not significant for the ICGC or Hammersmith methylation datasets and thus did not validate our TCGA methylation findings. Association between expression of ESRRA and TGFβR3 with overall survival in the ICGC and Tothill datasets was significant; however the association was protective and went in the opposite direction to the TCGA results. Therefore, the expression results in both datasets did not validate our TCGA expression findings.

Expression and methylation of ESR1, ESRRA as well as expression of TGFβR3 were determined in high grade serous ovarian cancer cell lines PEA1, PEA2, HEYA8, SKOV3 and IGROV1 as well as a control normal ovarian epithelium cell line. Interestingly both ESRRA and TGFβR3 shows a significantly higher level of expression (Figure 3A) in all high grade serous ovarian cancer cell lines than in OSE-C2 (though not significantly in the case of PEA1 for TGFβR3), unfortunately methylation did not show a concomitant decrease, in these lines however expression is transcriptionally regulated by greater number of epigenetic factors than methylation alone (Figure 3B). Pyrosequencing primers could not be designed to the sequence surrounding the TGFβR3 CpG site identified by the Illumina array probe so methylation could not be determined. Expression of ESR1 (Figure 3A) was very low in the OSE-C2 control cell line as well as in ovarian cancer cell lines PEA1, PEA2 and HEYA8 with raw Ct values of around 35. SKOV3 shows a significantly higher level of expression of ESR1 and IGROV1 as a hormone receptor-negative cell lines does not express ESR1 (with a raw Ct value of 40) as expected. Methylation of the selected ESR1 intragenic CpG site (Figure 3B) differs in the cancer cell lines with the changes not coinciding with the observed changes in expression. PEA1, PEA2 and HEYA8 show significantly increased altered methylation although not expression. SKOV3 and IGROV1 show significantly lower methylation than OSE-C2. Chemoresistant paired ovarian cancer cell lines were also tested for methylation and expression levels of the aforementioned genes, specifically A2780/CP70, PEO1/PEO4, PEA1/PEA2, and PEO14/PEO23.

**Figure 3.**
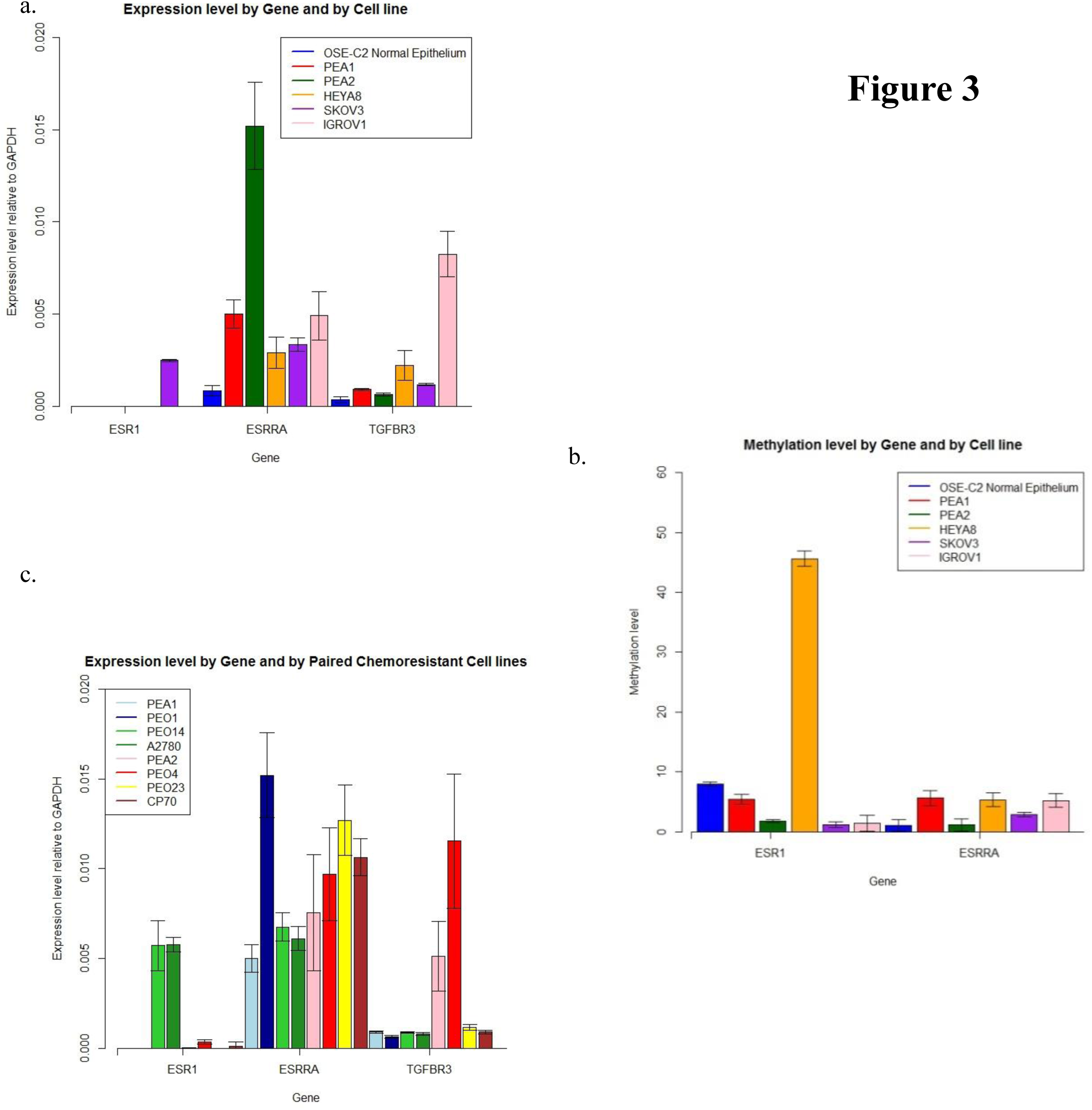
a. Expression level by Gene and by Cell line b. Methylation level by Gene and by Cell line c. Expression level by Gene and by Paired Chemoresistant Cell lines

PEO14 and PEO23 is the only cell line pair which shows a differential level of ESR1 expression (Figure 3C). PEA2 shows higher expression of ESRRA (Figure 3C) than PEA1, no significant difference is observed between other cell lines. The higher level of expression in PEO4 than PEO1 has a borderline significant p-value (p=0.054). TGFβR3 expression (Figure 3C) is significantly higher in PEA2 than PEA1 however no significant difference in expression exists between the other three pairs of cell lines. PEA2 shows a significantly more highly methylated ESRRA than PEA1 (Figure 3C). Methylation data is unavailable for the gene TGFβR3 or the Chemoresistant paired ovarian cancer cell lines PEO1, PEO4, PEO14, PEO23, A2780 or CP70.

## DISCUSSION

This work had several limitations, most importantly with the biostatistical validation. The expression and methylation medians for both TGFβR3 and ESRRA found in TCGA and used to dichotomise the probes and create four categories could not be used in the validation datasets as the TCGA values were lower than those found in the validation datasets (Figure 4). Essentially this resulted in only a few patients or none being below the TCGA median for TGFRB3 and ESRRA and the cox model being too unstable with too few observations to adequately produce results. We therefore had to use medians found in each individual validation dataset; thereby preventing a true validation. Furthermore, different expression assays were used in TCGA and ICGC again preventing a true validation.

**Figure 4.**
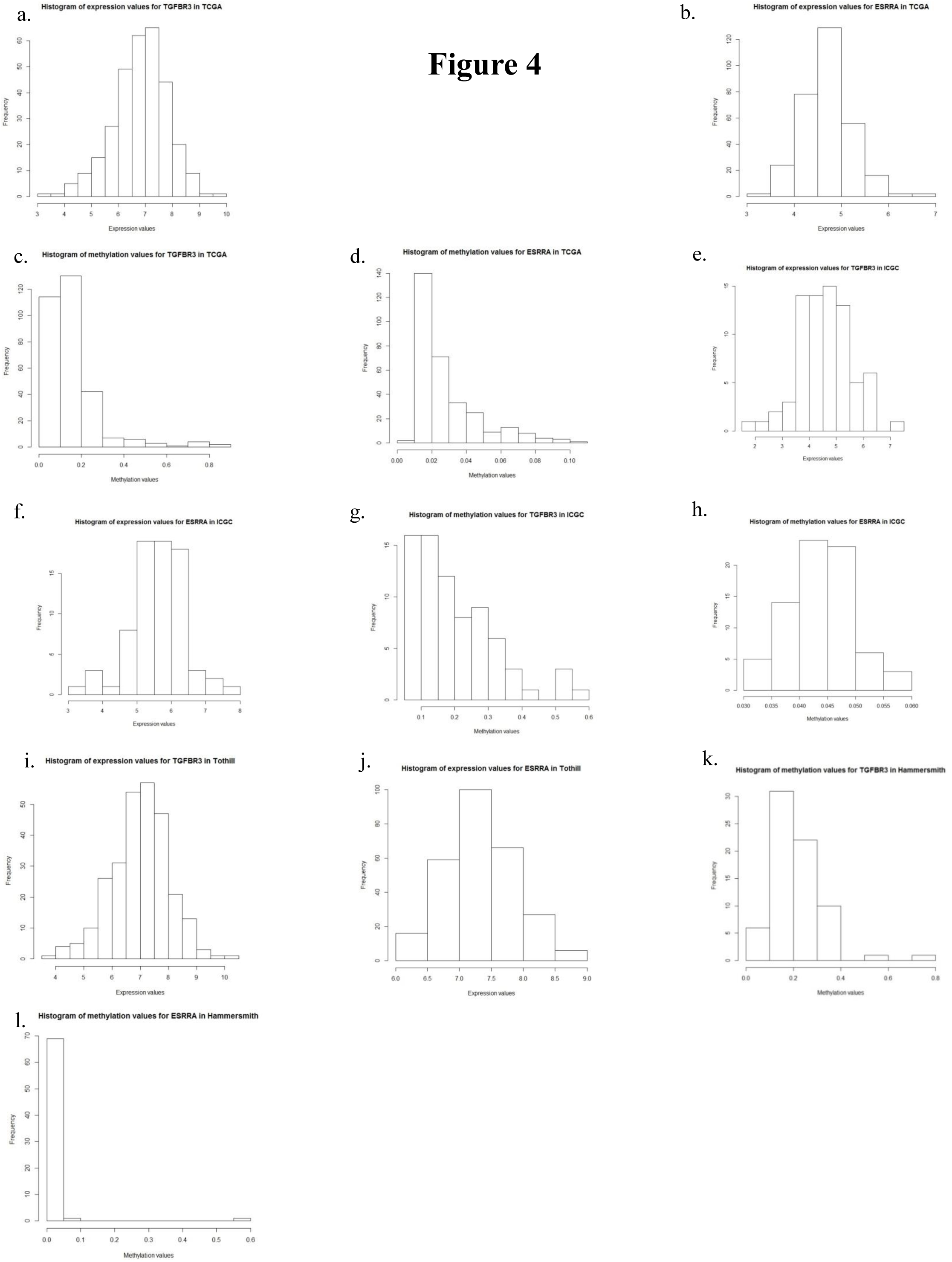
a. Histogram of expression values for TGFBR3 in TCGA b. Histogram of expression values for ESRRA in TCGA c. Histogram of methylation values for TGFBR3 in TCGA d. Histogram of methylation values for ESRRA in TCGA e. Histogram of expression values for TGFBR3 in ICGC f. Histogram of expression values for ESRRA in ICGC g. Histogram of methylation values for TGFBR3 in ICGC h. Histogram of methylation values for ESRRA in ICGC i. Histogram of expression values for TGFBR3 in Tothill j. Histogram of expression values for ESRRA in Tothill k. Histogram of methylation values for TGFBR3 in Hammersmith n. Histogram of methylation values for ESRRA in Hammersmith.

Despite the biostatistical validation, these genes have interesting biological functions potentially impacting ovarian cancer prognosis. Estrogen related receptor (ESRR) members are biologically active orphan nuclear receptors (11-12) which are closely related to the estrogen receptor (ER) family through the sharing of target genes, co-regulators and promoters. Additionally by targeting the same set of genes, the ESRRs seem to interfere with the ER-mediated estrogen response and estrogen signaling in various ways (11). ESRRA has been implicated in risk and poorer OS in ovarian cancer, with increased expression shown in both tumours and ovarian cancer cell lines (12). The authors theorised that in serous ovarian cancer cancer up-regulation of ESRRA may occur. ERRs although unable to bind estradiol, regulate the oestrogen response via protein-protein interactions with oestrogen receptors (ERs) (13), binding of oestrogen response elements (EREs) and recruitment of the same co-regulatory molecules as ERα/β (14), which regulate transcriptional programmes in response to oestrogen signalling. Oestrogen signalling via ERα/β activity has a role in regulating expression of genes involved in immune system function (15) including components of the TGFβ pathway (16), which in turn pertains to the maintenance of the inflammatory microenvironment and progression of cancer (14). More specifically, the transforming growth factor beta (TGFβ) pathway has long been of interest in carcinogenesis; in normal cells and in early stages of cancer development, TGFβ functions as a tumour suppressor. Meanwhile in higher stage cancerous cells, TGFβ functions as a tumour promoter, promoting progression and metastasis in later stages of cancer and possibly repressing TGFβR3 expression (17-19). In ovarian cancer, TGFβR3 is thought to function as a tumour suppressor, with it being implicated in invasiveness, metastasis, cell migration and motility and further showing downregulation in tumours and in ovarian cancer cell lines (18-19). TGFβR3 is the most well characterised receptor of the TGFβ pathway, signalling of which is documented as being oncogenic in late stage cancers (17).

The expression data for ESR1, ESRRA and TGFβR3 allowed us to identify a cell line in which to conduct functional studies on these putative serous ovarian cancer biomarkers. SKOV3 expresses all four genes, showing the only convincing level of ESR1 expression; therefore permitting interaction studies of these genes interact in serous ovarian cancer cells with oestrogen signalling and potentially affect cell survival. Future studies would firstly investigate the consequences of altered oestrogen availability on TGFβR3 expression as well as siRNA knock down of ESR1. Estradiol has been shown to regulate expression of components of the TGFβ pathway (16) and therefore potentially TGFβ3. The IGROV1 cell line is confirmed to be ESR1 negative therefore will provide a control cell line into which ESR1 can be introduced by an over-expressing plasmid to determine effect on TGFβR3 expression. ESRR genes are known to interfere and compete with ER signalling (14) and therefore may deregulate ER-mediated control of TGFβR3 expression. Therefore a siRNA knock-down of ESRRA will be utilised and effects on TGFβ3 expression assessed by RT-PCR and western blot. TGFβ has been shown to increase invasion in ovarian cancer via increased expression of matrix metalloproteinase (MMP) leading to matrix collagen cleavage and invasive potential (20). Expression of TGFβR3 will also be manipulated via siRNA and an over-expressing plasmid to determine what functional effect TGFβR3 over-expression has on the tumour cell to promote poor prognosis in HGSOC patients. An invasion assay could be performed in a 3D matrigel matrix using SKOV3 cells both untreated and treated with siRNA targeted to TGFβR3 and exposed to different TGFβ concentrations to determine whether elevation of TGFβR3 increases invasive potential in response to ligand binding. Effect on MMP expression could also be investigated. MTT cell proliferation assays could be employed to assess whether TGFβR3 expression has an effect on cell survival and caspase 3/7 assays to indicate the effect upon cellular apoptosis. The results indicate that expression and methylation of ESRRA and expression of TGFβR3 is higher in serous ovarian cancer tissue than normal tissue. This highlights the potential role of these genes in tumourigenesis leading to serous adenocarcinoma of the ovary. Aberrant expression of ESRRA in ovarian cancer has been noted previously (21). A limitation of the control cell line is that ovarian epithelium may not be the cell type from which the tumour originated, modern genetic and histopathological evidence suggest fallopian tube, endometrium and endocervical origins of epithelial ovarian cancer (22). Despite some significant differences in expression between chemosensitive and resistant cell lines, e.g. ESRRA in PEA1 and PEA2 the RT-PCR data for ESR1, ESRRA, and TGFβR3 does not indicate that altered expression of any of these genes may play a role in chemoresistance in ovarian cancer. The incompleteness of the methylation data is a limitation as a conclusion cannot be drawn regarding the potential for methylation of these genes as a biomarker for chemoresistance. The apparent absence of a trend in expression of these genes and resistance to platinum-based therapy suggests that TGFβR3 may not have an inhibitory role in apoptosis.

This paper is another key example of the importance of both biological and biostatistical validation in a prognostic biomarker study in adequate validation datasets and cell lines.

